# Proton FLASH preserves neurocognition across delivery techniques: implications for clinical translation in pediatric brain tumors

**DOI:** 10.64898/2026.05.29.728901

**Authors:** Devin Miles, Daniel Sforza, Nicole Tan, Yanbo Yang, Mina Akter, Xingyu Chen, Chanel Hutchison, Hawley Helmbrecht, Tyler Findlay, Lingshu Yin, Kan Ota, Masumi Umezawa, Yuncheng Zhong, Curtiland Deville, Matthew Ladra, Xun Jia, Charles Eberhart, Eric Raabe, Karin Walsh, Calixto-Hope Lucas, Heng Li, Lauren Jantzie, Robyn D. Gartrell, Sahaja Acharya

## Abstract

**Background:** Radiation therapy is integral to the curative treatment of childhood brain tumors but contributes to late neurocognitive impairment in survivors. FLASH (ultra-high dose rate, >40Gy/s) reduces normal-tissue toxicity in preclinical models, and proton-FLASH is currently the only modality capable of delivering ultra-high dose rates to the deep targets, such as pediatric brain tumors. However, two questions remain unresolved before clinical translation: (1) whether the FLASH effect can be achieved on synchrotron-based proton systems, which deliver protons in discrete spills that may be insufficient to cover a clinical target within a single delivery, and (2) which dose-rate metric, among the multiple definitions currently used in the field, best predicts the biological FLASH effect.

**Methods:** C57BL/6 mice (7-8 weeks) received 10 Gy whole-brain RT via a clinical Hitachi ProBEAT synchrotron with CBCT-guided delivery, using three transmission-beam techniques: single-spill pencil beam scanning (SS PBS), multi-spill PBS with ∼2-second inter-spot delay (MS PBS), and passive scatter (PS), compared to conventional (CONV) delivery and unirradiated controls (n=24–28/group, equal sex distribution). Dose rate was quantified using three frameworks: field dose rate (FDR), PBS dose rate (PBSDR), and dose-averaged dose rate (DADR). Recognition memory was assessed by novel object recognition (NOR) at 6 weeks post-RT, and cognitive flexibility was assessed via touchscreen visual discrimination and reversal learning at 14 weeks. Hippocampal neuroinflammation was evaluated by immunofluorescence and immunohistochemistry for Iba1, NeuN, and GFAP.

**Results:** FLASH conditions were met by SS PBS and PS under all three dose-rate definitions, but MS PBS qualified as FLASH only by DADR. Despite this, neuroprotection was preserved across all three FLASH techniques: discrimination index was significantly higher for SS PBS (P=0.021), MS PBS (P=0.008), and PS (P<0.001) versus CONV, with no significant difference between FLASH techniques. On touchscreen testing, FLASH-treated females demonstrated preserved cognitive flexibility (P=0.047 vs. CONV on reversal learning correct trials). Iba1^+^ microglia were reduced in FLASH compared to CONV mice, with morphology suggestive of preserved homeostatic state.

**Conclusions:** Synchrotron-based proton FLASH preserves neurocognitive function across all delivery techniques, including under multi-spill delivery essential for treating clinical-scale pediatric brain tumors. Critically, this neuroprotection was observed even for deliveries that qualified as FLASH only by DADR, identifying DADR as the dose-rate metric most relevant to the biological FLASH effect with direct implications for clinical trial design and dose-rate reporting standards.

## Introduction

Radiation therapy (RT) is a critical therapeutic modality for childhood tumors; however, its use in pediatric brain tumors is limited by toxicity, particularly neurocognitive impairment and radiation necrosis. Survivors of childhood brain tumors can exhibit significant neurocognitive impairment, which affects academic performance,^1^ social-emotional functioning,^2^ and the ability to live independently as an adult.^3^ Additionally, the risk of brainstem radiation necrosis^4,5^, which can lead to permanent neurologic injury or death, severely limits the role of dose-escalation for aggressive tumors close to the brainstem, such as medulloblastoma. Mitigating these toxicities while preserving tumor control is therefore a central unmet need in pediatric radiation oncology.

FLASH, a novel method of RT that uses ultra-high dose rates (UHDR, >40Gy/sec), has been shown to decrease toxicity compared to conventional RT (CONV, ≤2Gy/sec),^6^ known as the “FLASH effect.” The FLASH effect has been characterized in a preclinical setting with electrons^7^ and subsequently with photons^8^; however, currently, the only RT modality that can deliver FLASH to deep seated targets in humans is proton FLASH.^9^ The first-in-human proton FLASH trial treated painful extremity bone metastases in adults^9^ and a small number of preclinical studies have examined proton-FLASH in the brain with mixed results on neurocognitive protection.^10,11^ To date, no proton center has investigated FLASH in pediatric patients, and none has used a synchrotron-based system, which uniquely enables dynamic energy modulation of the proton beam without ridge filters or other passive attenuators. This allows precise conformal delivery, essential for sparing normal brain tissue in pediatric patients.

Two unresolved questions stand in the way of clinical translation of synchrotron-based proton-FLASH. First, can UHDR conditions be achieved with a synchrotron-based system at all? Prior proton-FLASH studies have primarily used cyclotron-based systems, which deliver protons as a continuous high-current beam. Synchrotrons, in contrast, deliver protons in discrete spills typically separated by ∼2 seconds, and the charge deliverable per spill is inherently limited. It is unknown whether sufficient charge can be extracted within a single spill to achieve UHDR, and whether the FLASH effect is preserved when larger targets require multi-spill delivery. This question has direct clinical relevance because the volumes treated in pediatric brain tumors, particularly under craniospinal irradiation, will require multi-spill delivery on currently available systems.

Second, which dose-rate metric best predicts the biological FLASH effect? Field dose rate (FDR), pencil-beam scanning dose rate (PBSDR),^12^ and dose-averaged dose rate (DADR)^13^ yield substantially different values for the same delivery, particularly for multi-spill PBS plans where inter-spot temporal gaps are large. This dosimetric inconsistency poses a direct challenge to FLASH trial design and clinical implementation, because the same delivery may or may not qualify as FLASH depending on which metric is used. To date, no study has paired these dosimetric definitions with biological endpoints to resolve which metric tracks the FLASH effect.

Here, we address both questions through the development and initial application of synchrotron-generated image-guided proton FLASH for murine whole-brain RT. We hypothesize that the FLASH effect, defined as reduced neurocognitive impairment compared to CONV, can be achieved under one or more synchrotron delivery conditions: (1) single-spill (SS) PBS, where the entire field is delivered within one beam spill; (2) multi-spill (MS) PBS, where each beam spot is delivered in a separate spill ∼2 seconds apart; and (3) passive scatter (PS), where the field is irradiated uniformly with a single-scattered beam and collimation. By pairing rigorous dosimetric characterization (FDR, PBSDR, DADR) with neurocognitive readouts (NOR, touchscreen visual discrimination, and reversal learning), neuroimaging, and microglial analysis, we test whether the FLASH effect is preserved across these synchrotron-specific delivery modalities and identify which dose-rate metric tracks the neurocognitive biological response.

## Methods

### Beam Specification

Irradiations at CONV and FLASH dose rates were performed using 142.4 MeV protons, delivered using a fixed horizontal proton beamline supplied by a clinical Hitachi ProBEAT synchrotron (**Fig. 1A**). Details on the initial characterization and commissioning of this beam have been published.^14^ Briefly, to facilitate FLASH irradiation, the radiofrequency (RF) extraction power was increased to achieve a complete spill of all deliverable protons in a short time frame, typically under 200 milliseconds. Due to the mechanism by which protons are produced and accelerated in a synchrotron, the maximum deliverable dose in FLASH mode is presently assumed to be limited by the maximum charge per spill, roughly 5-6 nC in our system, where an additional 2-second delay is required to replenish the charge before another spill can be performed. The clinical beam monitors in the nozzle were disabled for FLASH irradiation and replaced with a FLASH-ready transmission strip ionization chamber (IC-64, PTC) and 10 kHz multi-channel electrometer (I-128, PTC). Fast readout from the electrometer is converted into a simulated square pulse and fed back into the synchrotron controller to achieve fast control of the beam in FLASH mode. Absolute dose verification is conducted for all experiments before animal irradiations using a large-area Faraday cup (PTC), which is used to cross-calibrate the strip ionization chamber.

**Figure 1.**
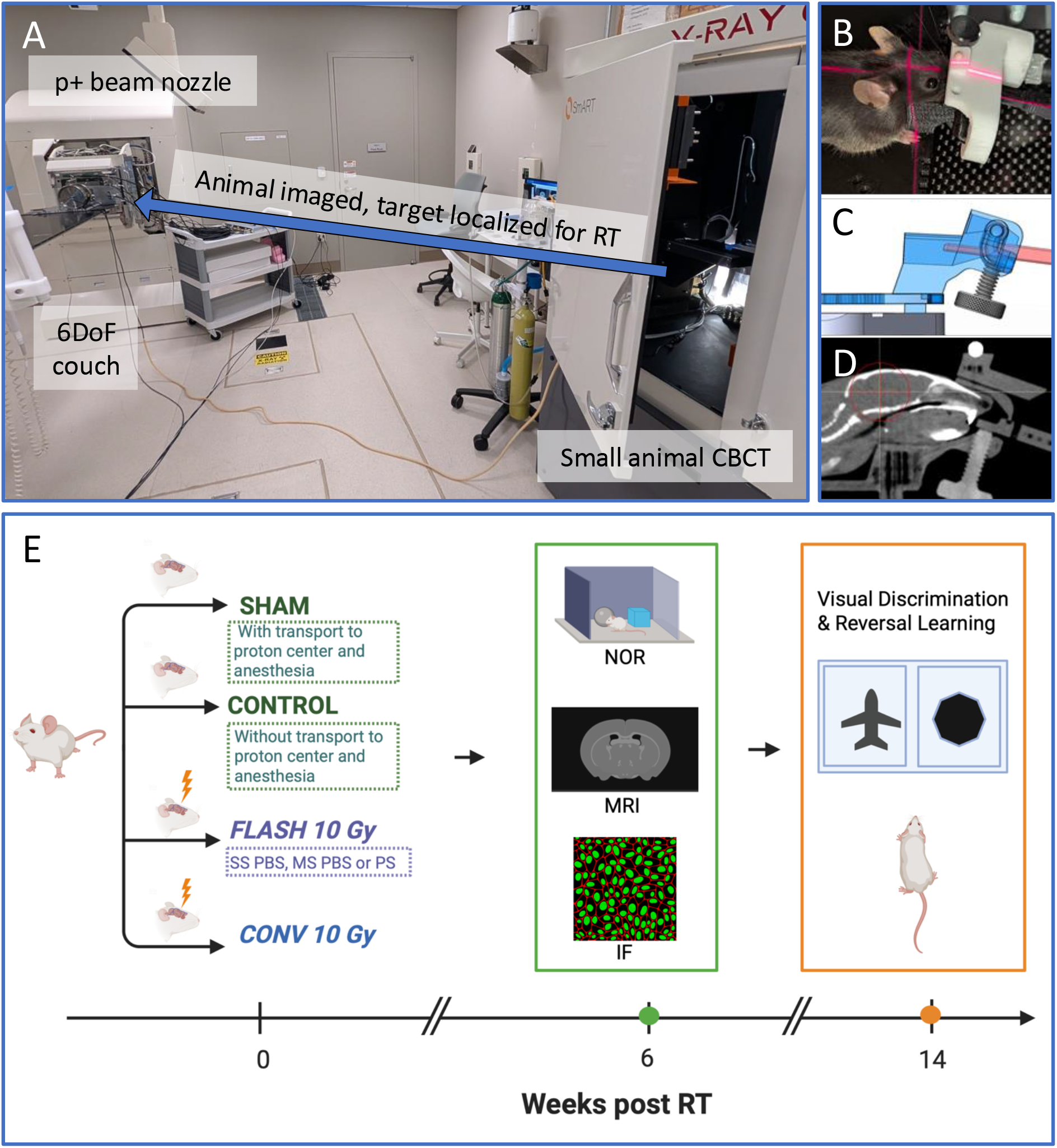
Overview of mouse IGRT setup and experimental design. (A) Animals were imaged using a small animal CBCT cabinet and manually moved to a 6-degree-of-freedom treatment couch in the horizontal proton beamline. (B-C) Reproducible mouse setup utilized rigid cranial fixation via a 3D printed immobilization tool with a locking bite-bar, with isoflurane input and indexed mounting fixture between the CBCT and proton couch. (D) The treatment isocenter was localized in 3D relative to the bb fiducial marker embedded in the nose cone. The bb was then set up to the proton isocenter, and a shift to the target isocenter was applied. (E) Experimental timeline from irradiation using varying delivery techniques at day 0, followed by NOR, MRI, and IF at 6-weeks post-irradiation. Visual discrimination and reversal learning assessments were performed starting at 14 weeks post RT.

### Treatment field specifications

A commissioned model of our 142.4 MeV beam in TOPAS was used to design radiation fields to achieve an average physical dose of 10 Gy to the whole mouse brain in a 1 × 1-cm^2 square beam using a ‘shoot-through’ technique with the entry plateau region of the proton Bragg peak.

To assess the impact of pencil-beam scanning (PBS) and instantaneous dose rates on all assessed endpoints, proton beams were designed using three delivery paradigms: (1) SS PBS, (2) MS PBS, and (3) PS. In SS and MS delivery, the mouse brain is treated using the same square 3-by-3 spot pattern via PBS. Here, SS is considered the ‘default’ paradigm where an entire target may be treated to its full dose within a single beam spill in FLASH mode (i.e., high/variable local dose rate, high average dose rate). A ‘peaked’ RF extraction pulse was optimized for SS delivery to achieve the highest possible deliverable charge per spill.^15^ This results in a higher beam current at the beginning of a spill, and a decrease in current (thus, dose rate) toward the end of the beam spill, despite the charge per spot being held constant. MS delivery was considered to emulate a ‘worst-case’ in synchrotron charge limitations where a target may not be treatable in its entirety within a single spill (i.e., high/variable local dose rate, low average dose rate). This is especially important when moving from small targets in mice to larger targets in humans as targets in humans will require MS delivery. The single-scattered beam was considered as a way to ensure high local and volumetric dose rate independent of beam scanning (i.e., high/uniform local dose rate, high average dose rate). To facilitate PS, a simple conical single scatterer and square collimator were optimized in TOPAS and fabricated using readily available Cerrobend in our clinic and cast using 3D-printed molds.

### Image-guided animal irradiation workflow

All animal experiments were performed in accordance with national guidelines and approved by the Johns Hopkins University School of Medicine Animal Care and Use Committee. Healthy 7-8-week old C57BL/6 mice (Jackson Labs) were transported to our proton therapy center from a central housing facility on the morning of each irradiation day and returned to housing on the same day. Male and female mice were used equally for most experiments. Only female mice were kept for touchscreen analysis. Mice received FLASH or CONV WBRT as described below, and SHAM mice were transferred to the proton facility and received anesthesia similar to the treated mice without undergoing RT.

Mice were immobilized for imaging and irradiation using a custom-designed 3D printed tool, featuring rigid cranial immobilization mediated by a bite block with a built-in anesthesia nosecone and bb fiducial marker (**Fig. 1B-C**). The immobilization device also includes a baseplate designed for rigid indexing onto the irradiation and imaging couches. Once positioned in the immobilization, mice were imaged using a small-animal cone-beam CT (PXI SmART), which was used to identify the treatment isocenter per experiment and the isocenter’s displacement from the bb fiducial marker (**Fig. 1D**). The immobilized animals were then transferred onto the proton treatment couch, where the fiducial marker was aligned to room lasers and a shift applied to align the target isocenter to the beamline. Animal setup reproducibility was quantified by co-registering CTs of immobilized animals in Raystation v2023B, focusing on rigid alignment of the immobilization device and fiducial markers. The Dice similarity coefficient between brain regions of interest and the distance-to-agreement between bony features of the skull were calculated (n = 10 CTs assessed).

An end-to-end test was performed to quantify the uncertainty associated with our image-guided RT workflow. A 3D-printed film holder including a centered bb fiducial marker was designed to fit onto the mouse immobilization device. EBT3 film was laser-cut to ensure a highly reproducible fit onto the film phantom, including a cutout for the bb fiducial, which was considered the reference marker target for the central axis of the proton beam. The immobilization device with film was imaged using CBCT and placed into the beam, similar to mouse irradiations. E2E irradiations were performed using the complete image-guided workflow (CBCT-guided shift from fiducial). Irradiated films were scanned and imported into Matlab for analysis. An in-house script was written to calculate the displacement between the film fiducial cutout and the center of the irradiated area. The maximum displacement between the fiducial marker and the radiation field center was considered to be our overall uncertainty for beam delivery. Dose and dose rates were recalculated considering this setup uncertainty to quantify the robustness of our radiation delivery.

### Whole-brain irradiation

Immediately prior to irradiation, mice were individually anesthetized using gaseous isoflurane supplied by room air (Kent Scientific) and placed onto the immobilization tool for imaging and irradiation where they continued to receive anesthesia throughout imaging and RT. Mice received a single fraction of 10 Gy to the whole brain in SS, MS, or PS delivery, in FLASH or CONV dose rate modes (n=12-14/treatment group/sex). A delivered dose of 10 Gy was chosen for comparison with prior published experiments describing the FLASH effect in mouse brain irradiation.^6^ Proton relative biological effectiveness was not factored into the delivered dose because the Bragg peak was not utilized. The delivered dose and beam time structure was recorded for each mouse irradiated at FLASH dose rates using the transmission strip ion chamber. A SHAM irradiation arm (n=10-12/sex) was included, where un-irradiated mice received isoflurane within a week of the radiation cohorts. Following irradiation, mice were transported to a central housing facility where animals were monitored by housing staff daily, and weights were tracked weekly. A control arm (n=6/sex) was also included, which represented mice that did not undergo irradiation, anesthesia exposure or transport to the proton facility.

Local dose rates were calculated for FLASH delivery using the measured delivery time structure and the calculated dose per beam spot in a water phantom from the TOPAS model. Local dose rate was calculated using three metrics: (1) field dose rate, calculated as the local dose divided by the total beam delivery time, (2) PBS dose rate,^12^ (3) dose-averaged dose rate.^13^ Dose-rate volume histograms were calculated for each dose rate definition using a representative brain region of interest (ROI) superimposed in the dose rate maps. Robustness in the delivered doses and dose rates against setup and targeting errors were assessed via misalignments of the brain ROI and dose rate maps, based on the results of the E2E test described in the previous section.

### Novel object recognition

Short-term memory is commonly impaired in children who have undergone cranial radiation.^16^ To measure memory deficits in mice, we used novel object recognition (NOR) testing, which is a well-established murine assay for characterizing recognition memory and has previously been used in multiple FLASH studies.^6,17,18^ All mice underwent NOR at 6-weeks post RT. The test consists of three sequential phases: habituation, familiarization and recognition. During habituation, mice were acclimated to a quiet, dimly lit testing arena for three days. Each day, they were placed in an empty testing box and allowed to freely explore for 10 minutes before being returned to their housing. On the fourth day, mice underwent the familiarization phase, during which they interacted with two identical objects taped 16 cm apart in opposite corners of the box. Mice were allowed to explore both objects for 5 minutes. An interaction was defined as the mouse having its nose within 2 cm of an object. Interactions were recorded using a video tracking system (ANY-maze, Stoelting Co.), which quantified both the number of interactions and the total time spent investigating each object. Following a 5-minute inter-trial interval in the home cage, one of the familiar objects was replaced with a novel object, initiating the recognition phase. Mice were returned to the arena and allowed to explore for an additional 5 minutes. The positions of the familiar and novel objects were counterbalanced across animals to prevent spatial bias. Interaction times with each object were recorded, and recognition memory was quantified using the discrimination index (DI), calculated as:

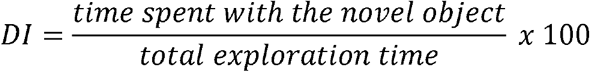

Mice that spent ≤ 7 seconds interacting with either object in either familiarization or recognition phases were re-tested on the following day. Re-testing consisted of repeating the familiarization and recognition phases. Mice that spent ≤ 1 second interacting with either object during re-testing were excluded from the analysis as these mice were likely not spending enough time with either object to appropriately assess recognition memory.

### Visual discrimination and reversal learning touchscreen testing

Processing speed is another commonly impaired domain in children who have undergone cranial radiation.^9^ This testing platform is analogous to Cambridge Neuropsychological Test Automated Battery (CANTAB), a widely validated, digital, touch-screen cognitive assessment tool and platform commonly used in human neuropsychological assessments.^19^ Processing speed and cognitive flexibility can be measured through tests of visual discrimination and reversal learning. Twenty-seven female mice (6 SS FLASH, 11 CONV, 10 SHAM) underwent touchscreen testing. Touchscreen testing was performed in female mice to evaluate cognitive flexibility in the sex group that did not demonstrate a significant FLASH benefit on NOR, allowing assessment of whether more sensitive paradigms could detect a treatment effect. The smaller number of FLASH-treated females in the touchscreen cohort (n=6) relative to CONV (n=11) and SHAM (n=10) reflects relocation of testing chambers to a different facility before all FLASH animals could complete the protocol.

Testing was conducted in light- and sound-attenuating chambers equipped with a touchscreen apparatus (Conclusive Solutions, Sawbridgeworth, UK). The touchscreen, divided into two response windows and covered with an acrylic plate, was controlled for stimulus presentation and response tracking. Correct responses were rewarded with 20 mg food pellets, delivered via a dispenser and signaled by light and sound cues. Mice were weight restricted to 85% of their free feeding weight to ensure that food remained a strong reward.

Following attainment of weight criteria, mice were acclimated to the operant chamber and trained to retrieve food rewards from the magazine. 10 rewards were placed in the magazine and mice that retrieved all 10 rewards within 30 min advanced to lever-press training, during which rewards were earned exclusively via responses on an ultrasensitive lever and paired with an auditory tone and magazine illumination. Advancement to visual discrimination testing required collection of 30 rewards within 30 min.

Visual discrimination and reversal testing were conducted as previously described.^20,21^ In brief, during visual discrimination, mice were trained to discriminate between two novel, approximately equally luminant stimuli presented pseudorandomly across 30-trial sessions using nose-poke responses, with correct stimuli counterbalanced across mice and treatments.

Correct responses were rewarded, whereas incorrect responses resulted in a timeout; correction trials followed errors. Once mice achieved criterion for visual discrimination, they proceeded to reversal learning where stimulus–reward contingencies reversed. Visual discrimination and reversal learning criteria were defined as ≥24 correct responses per 30 trials across two consecutive sessions.

For visual discrimination and reversal learning phases, performance metrics were aggregated across sessions for each mouse. The primary outcome measures included: days to graduation (number of days from first session to achieving graduation criteria), mean correct trials per session, mean completed trials per session, mean response latency for correct trials, mean response latency for incorrect trials, mean total response latency, and mean magazine latency.

Outliers were identified to exclude trials or sessions with off-task behavior or equipment issues prior to calculating outcome measures. For trial counts (completed trials and correct trials), performance was expected to improve over the course of training. To account for this learning effect, a linear trend was fitted to each mouse’s data, and z-scores were calculated from the residuals. Sessions with z-scores exceeding ±2.576 were flagged as outliers. If the fitted slope was negative (contrary to expected improvement), residuals were calculated from the mean rather than the regression line, as a negative trend likely reflects inconsistent performance rather than true learning. The first five sessions of each phase were excluded from outlier detection to allow for initial task acquisition. For response latency measures (correct, incorrect, and total), z-score-based outlier detection similarly accounted for expected decreasing trends over time. However, sessions with latencies of 15 seconds or less were retained regardless of z-score, as these values reflect appropriate task engagement. For magazine latency, a fixed threshold was applied, sessions in which the mouse took 15 seconds or longer to retrieve the reward were excluded, as prolonged magazine latencies indicate disengagement of the reward retrieval.

### MR imaging at 6 weeks post RT

Nine female mice (n=3/treatment group) underwent brain MRI (Bruker 7T simultaneous MRI/S-PET scanner) at 6 weeks post RT before being sacrificed for histopathology, immunofluorescence (IF) and immunohistochemistry (IHC). The following whole brain MRI sequences were acquired: T1 with and without contrast, T2 and diffusion weighted imaging. The following regions of interest (ROIs) were segmented: ventricles, hippocampus, corpus callosum, external capsule, thalamus, cortex, basal ganglia. Hippocampus, corpus callosum and supratentorial white matter pathways are critical for memory and processing speed and vulnerable to RT mediated injury in pediatric brain tumor survivors.^22,23^ Regions of T1 contrast enhancement concerning for necrosis and regions of T2 hyperintensity concerning for edema were also segmented. Mean ADC of the hippocampus, corpus callosum and external capsule were also calculated.

### Immunofluorescence (IF) staining, image processing and analysis

Six female mice (n=2/treatment group [CONV, SS FLASH, SHAM]) were euthanized via cardiac perfusion of saline 6 weeks after RT. Brains were cut sagittally into two equal hemispheres and the left hemisphere was placed in 4% paraformaldehyde (PFA) for 24 h. Brain hemispheres were taken from PFA and cut coronally into three - 4 mm slices and then brought to histology core to prepare formalin-fixed paraffin-embedded (FFPE) blocks with all three slices included in each block. FFPE blocks were then cut into 5 μm immunoblank slides. For every 10 slides cut there was one stained with hematoxylin and eosin (HE). HE slides were reviewed with a neuropathologist (C-HL) to identify sections with the ROI including hippocampus and white matter.

IF staining was performed using CoraLite® Plus 488-conjugated GFAP antibody (1:200, CL488-16825, Proteintech), CoraLite® 555-conjugated NeuN antibody (1:200, CL555-26975, Proteintech), CoraLite® Plus 647-conjugated IBA1 antibody (1:100, CL647-81728, Proteintech) and DAPI(1:1000, 10236276001, Roche). IF stained slides were examined using the Leica Mica and imaged at 20X magnification using the widefield fluorescence mode. Background was removed using the THUNDER imaging computational clearing on the Leica Mica Microhub.

Representative images within the hippocampus from one mouse in each group (SHAM, CONV, FLASH) were chosen, and each channel was optimized for best image visualization (Fig 6A). Images were imported and annotated in QuPath.^24^ These images were confirmed to be representative of ROI by neuropathologist (C-HL), including images within hippocampus and white matter. Within the tissue annotation, nuclei were detected using the watershed cell detection algorithm on the DAPI channel, with cell expansion used to estimate whole-cell regions. Detected cells were classified using intensity-based thresholds from marker-specific measurements. Cells were considered GFAP-positive when cytoplasmic GFAP mean intensity met the defined threshold and cytoplasmic GFAP maximum intensity exceeded the upper threshold. NeuN positivity was assigned using nuclear NeuN mean intensity, and Iba1 positivity was assigned using cytoplasmic Iba1 mean intensity. Once analyzed, our neuropathologist was consulted to evaluate and help to confirm cells considered positive in a blinded fashion.^25^

### IBA1 immunohistochemistry staining, image processing and analysis

Representative images within the hippocampus from one mouse were chosen and Iba1 immunostaining was quantified in QuPath using manually defined areas of interest for each slide. Slides were analyzed after applying consistent H-DAB stain vectors across the dataset. Within each annotated region, color deconvolution was used to separate the DAB signal from hematoxylin, and Iba1-positive staining was quantified by thresholding the DAB optical density channel. Positive area analysis was performed at a requested pixel size of 1.0 µm. Tissue was identified using an optical-density-based tissue mask to exclude glass/background, and DAB-positive pixels were defined as pixels with DAB optical density ≥ 0.20. For each slide, Iba1-positive area was calculated as:

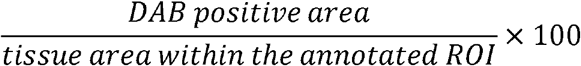

### Statistical Analysis

For NOR and touchscreen testing, groupwise comparison was performed using Wilcoxon rank-sum tests and for IF and IHC data, group mean bar plots were generated using stats packages in R (version 4.5.1).

## Results

### Radiation delivery summary

A summary of representative 2D dose maps, dose-rate maps, and radiation delivery metrics is shown in **Fig. 2**. Average doses measured at the field central axis were 10.04 ± 0.16 Gy (SS; **Fig. 2A**), 10.24 ± 0.20 Gy (MS; **Fig. 2B**), 10.01 ± 0.09 Gy (PS; **Fig. 2C**), and 9.92 ± 0.19 Gy (CONV). SS FLASH irradiation was delivered in 128.81 ± 84.98 ms on average (**Fig.2D**), whereas MS FLASH delivery required 16.4 ± 2.9s (**Fig. 2E**), despite comparable typical charge per beam spot and delivery time per spot between SS and MS modes. Notably, charge per spot and spot duration were more consistent in MS mode because protons were delivered only during the “peak” of the RF extraction signal. PS delivery was completed in the shortest overall delivery time at 76.87 ± 12.01 ms **Fig. 2F**), given a lack of the requirement for beam scanning. However, the average delivery time per beam spot for PS was approximately 10-fold greater than SS and MS. In CONV dose-rate mode, the full dose was delivered in 11.3 ± 0.6 s, which was faster than MS FLASH delivery (**Fig. 2Q**).

**Figure 2.**
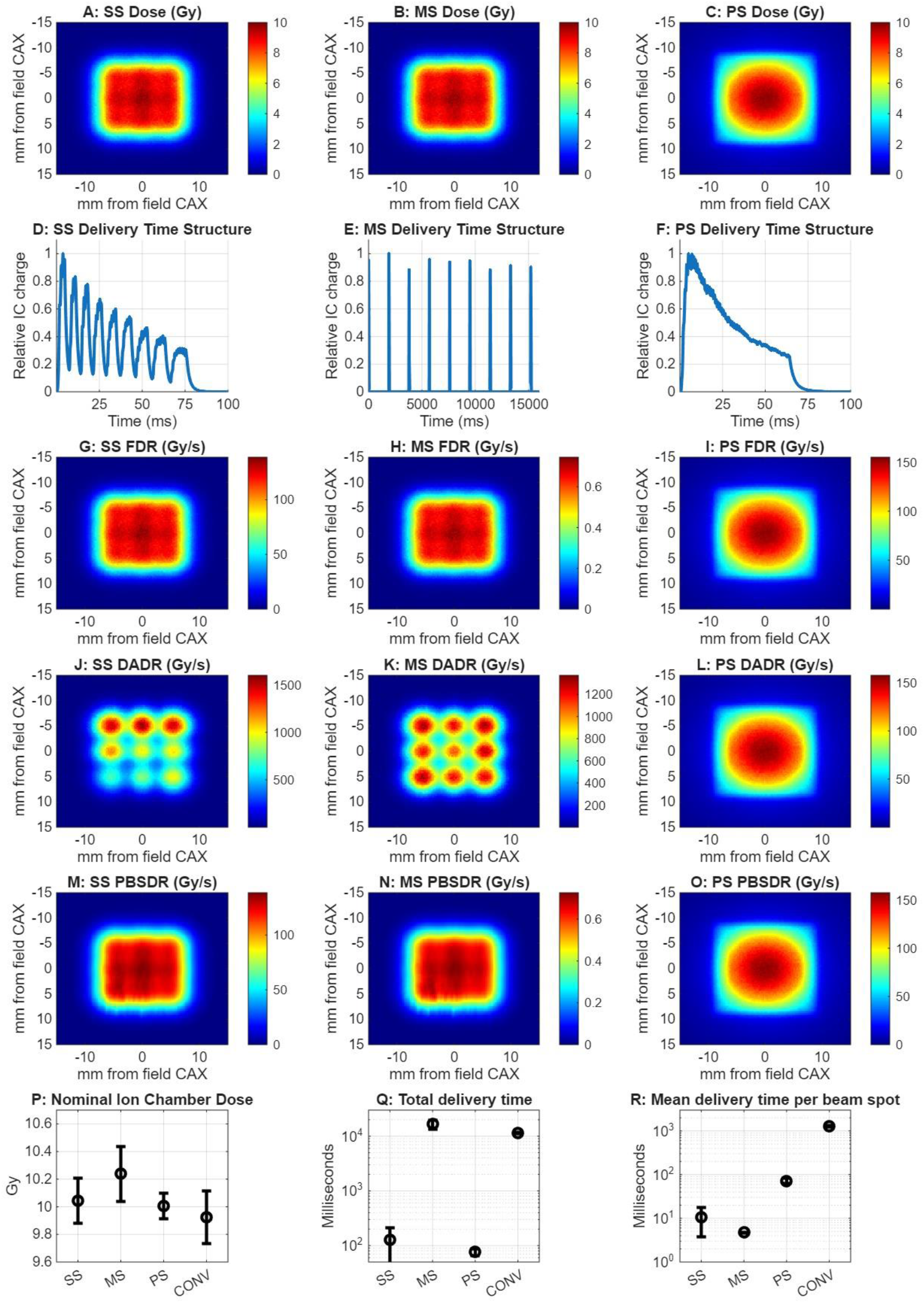
Representative dose and dose rates for UHDR radiation delivery (Single spill: SS, Multi-spill: MS, Passive scatter: PS, Conventional: CONV). (A-C) Dose maps for SS, MS, PS calculated in TOPAS. SS, MS, and CONV are identical in the simulation where time resolution is not considered, i.e. the same 3 × 3 spot placement is used. (D-F) Measured radiation delivery times, sampled using the transmission strip ion chamber with sampling rates up to 10 kHz. The synchrotron delivers UHDR using a ‘peaked’ RF extraction, where the initial beam current is high and decreases slightly throughout the beam spill. This peaked behavior is less apparent for MS than SS and PS delivery. (G-I) Field dose rate (FDR) maps per UHDR condition. (J-L) Dose-averaged dose rate (DADR) maps per UHDR condition. (M-O) Pencil beam scanning dose rate (PBSDR) maps per condition. (P) Nominal CAX doses as measured with the transmission ion chamber were consistent to within 2% across all delivery modes for animal irradiations in this manuscript. (Q-R) The total delivery time for all delivery modes and the mean delivery time per spot. Note that SS and PS achieve UHDR conditions in all dose rate definitions, while MS conditions only meet UHDR as defined by DADR. For plots with error bars, data is presented as the mean ± the standard deviation of all measured data per group.

Using the definitions of field dose rate (FDR; **Fig. 2G–I**) and PBS dose rate (PBSDR; **Fig. 2M–O**), UHDR conditions were achieved only for SS and PS delivery. Typical FDR values for MS delivery were below 0.6 Gy/s, while CONV achieved 1.6 Gy/s. In contrast, when dose-averaged dose rates (DADR) were calculated, UHDR was achieved for SS, MS, and PS delivery techniques. SS DADR was highly heterogeneous due to the peaked RF extraction (with higher DADR for initial spots than for later spots within the spill), whereas MS DADR was more consistent. A summary of dose-delivery metrics across SS FLASH, MS FLASH, PS FLASH, and CONV PBS deliveries is provided. All delivery techniques were within 3% of the intended 10 Gy nominal delivered dose (**Fig. 2P**), while total delivery time (**Fig. 2Q**) and mean delivery time per beam spot (**Fig. 2R**) varied substantially across techniques.

Targeting uncertainties from end-to-end film testing averaged 0.78 ± 0.15 mm in the superior– inferior direction and 0.65 ± 0.25 mm in the anterior–posterior direction. Summed in quadrature, the overall IGRT targeting uncertainty was 1.02 ± 0.29 mm. For robustness calculations, a 1 mm targeting uncertainty was assumed. The high Dice similarity coefficient for brain ROIs (0.85 ± 0.06) and low distance-to-agreement for bony anatomy (DTA; 0.37 ± 0.01 mm) across 10 representative animals indicated minimal inter-animal setup variability and were therefore not incorporated into the setup uncertainty estimate.

### Mouse whole-brain radiation dosimetry

Beam delivery for each mouse irradiated in UHDR mode was recorded with the transmission strip ionization chamber and used to compute delivered local dose rate, as well as dose- and dose-rate volume histograms (**Fig. 3A–D**). A wide range of SS dose rates was observed, reflecting variable delivery times during the early stages of the experiment; mice treated at a nominal dose rate below 40 Gy/s were excluded from this experiment and subsequent endpoints.

**Figure 3.**
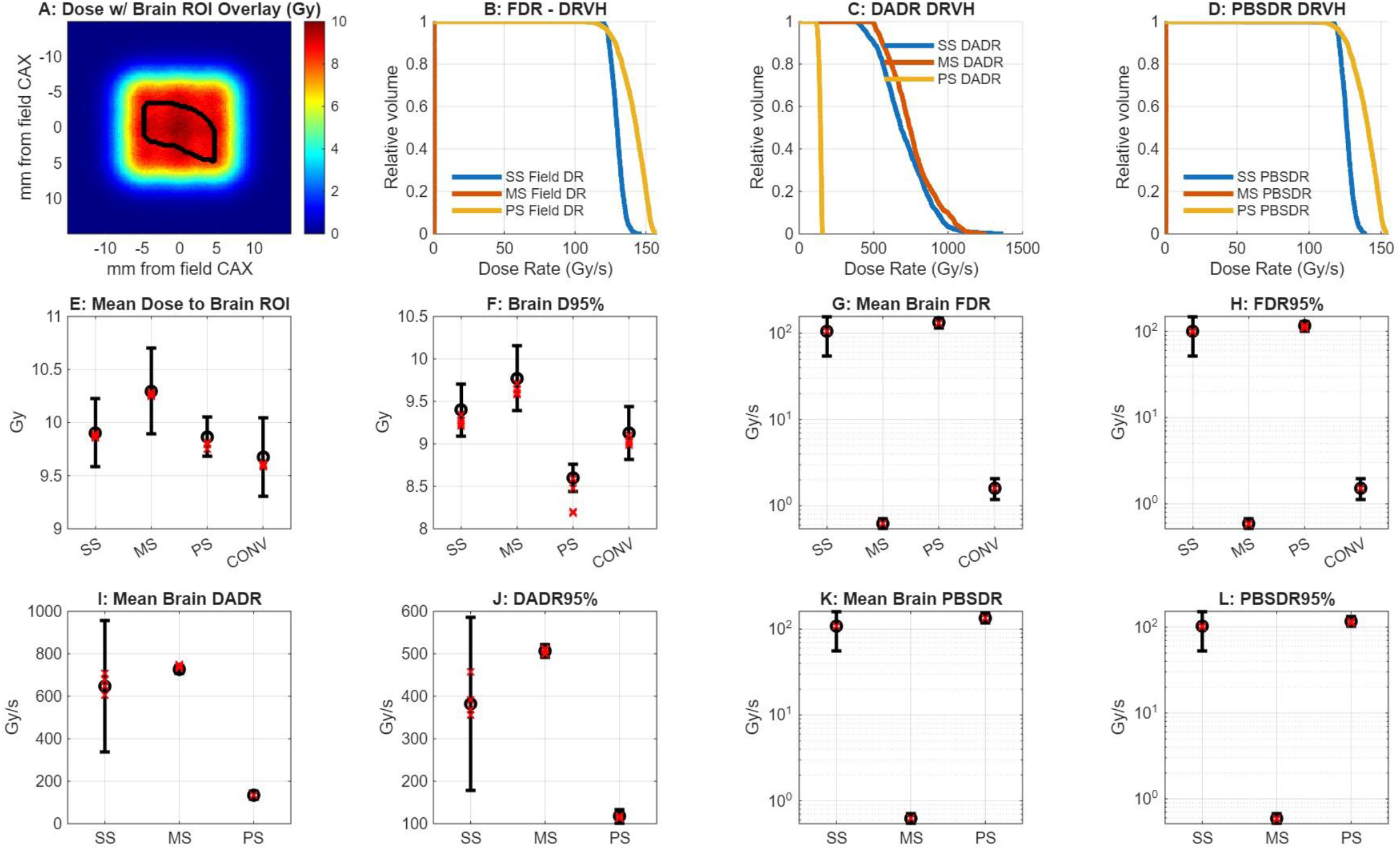
Brain DVH and DRVH metrics for all assessed delivery modes. (A) Representative mouse brain ROI overlaid on SS dose map. (B-D) Representative brain DRVH plots for all UHDR delivery modes using FDR, DADR, PBSDR dose rate definitions. Note that MS does not reach UHDR for FDR and PBSDR definitions. (E-F): Mean and D95% doses to the brain across delivery modes. Note that the mean doses are very consistent for all modes, but the D95% for PS is lower due to the lower uniformity of the beam. (G-L) Mean and DR95% dose rates to the brain ROI for all delivery modes, varying dose rate definition. Black markers are the mean ± standard deviation as measured for all mouse irradiations. Red markers are the mean for robustness scenarios where the ROI is offset by 1mm in each cardinal direction.

The mean dose delivered to the whole brain was 9.9 ± 0.32, 10.30 ± 0.40, 9.87 ± 0.18, and 9.67 ± 0.37 Gy for SS, MS, PS, and CONV deliveries, respectively (**Fig. 3E**). The minimum brain dose (D95%) was 9.4 ± 0.31, 9.77 ± 0.38, 8.59 ± 0.16, and 9.13 ± 0.31 Gy for SS, MS, PS, and CONV, respectively (**Fig. 3F**). PS delivery produced the lowest minimum brain dose across conditions, consistent with the greater heterogeneity of the SS system, although the mean brain dose was not significantly different from the other conditions. Brain D95% was the only metric that changed significantly under the modeled setup and targeting uncertainty (**Fig. 3F**), reaching a minimum of 8.18 Gy for a 1 mm anterior shift.

Using the FDR definition, UHDR conditions were achieved only for SS and PS. Mean FDR to the brain was 105.60 ± 51.36 Gy/s for SS, 0.63 ± 0.08 Gy/s for MS, 132.47 ± 17.34 Gy/s for PS, and 1.61 ± 0.43 Gy/s for CONV dose-rate conditions (**Figs 3G-H**).

When local dose rate was defined by DADR, UHDR dose rates to the whole brain were achieved for each FLASH delivery condition (**Fig. 3I–J**). DADR values were substantially higher for SS and MS than for the other dose-rate definitions, reflecting the short delivery times per spot and omission of the delays between spots. Mean DADR was 646.76 ± 310.38 Gy/s for SS, 725.77 ± 19.07 Gy/s for MS, and 134.68 ± 17.84 Gy/s for PS. DADR was not calculated for CONV delivery.

Using the PBSDR definition, UHDR conditions were again achieved only for SS and PS. Mean PBSDR was 108.07 ± 53.02 Gy/s for SS, 0.63 ± 0.08 Gy/s for MS, and 134.65 ± 17.84 Gy/s for PS (**Fig. 3K–L**). PBSDR was not calculated for CONV delivery.

### NOR testing at 6 weeks post RT

All mice underwent novel object recognition (NOR) testing at 6 weeks following irradiation and/or anesthesia exposure at the proton center, corresponding to an age of 13 weeks. Unirradiated control animals who were not exposed to anesthesia were tested at the same age. The final analyzed cohort comprised: CONV (n=20), SS (n=27), MS (n=23), PS (n=27), SHAM (n=21), and control (n=12).

A significant difference in recognition memory, as indexed by the discrimination index (DI), was observed across treatment groups (p=0.006). Pairwise comparisons revealed that mice treated with each of the three FLASH delivery techniques — SS, MS, and PS — demonstrated significantly higher DI values, reflecting superior recognition memory, compared to CONV-irradiated animals (CONV vs. SS: p=0.021; CONV vs. MS: p=0.008; CONV vs. PS<0.001; **Fig. 4A**). DI did not differ when comparing CONV (median, [IQR]: 50.79, [15.67]) to SHAM (50.62, [19.52]) or unirradiated control animals (59.15, [23.30]; all p>0.05). No significant differences in DI were observed among the three FLASH delivery techniques (all p>0.05). PS-irradiated mice demonstrated significantly higher DI compared to SHAM animals (p=0.009), though this difference did not reach significance relative to unirradiated controls (p=0.111). No other FLASH technique differed significantly from SHAM or control, and no significant difference was observed between SHAM and control groups (all p>0.05).

**Figure 4.**
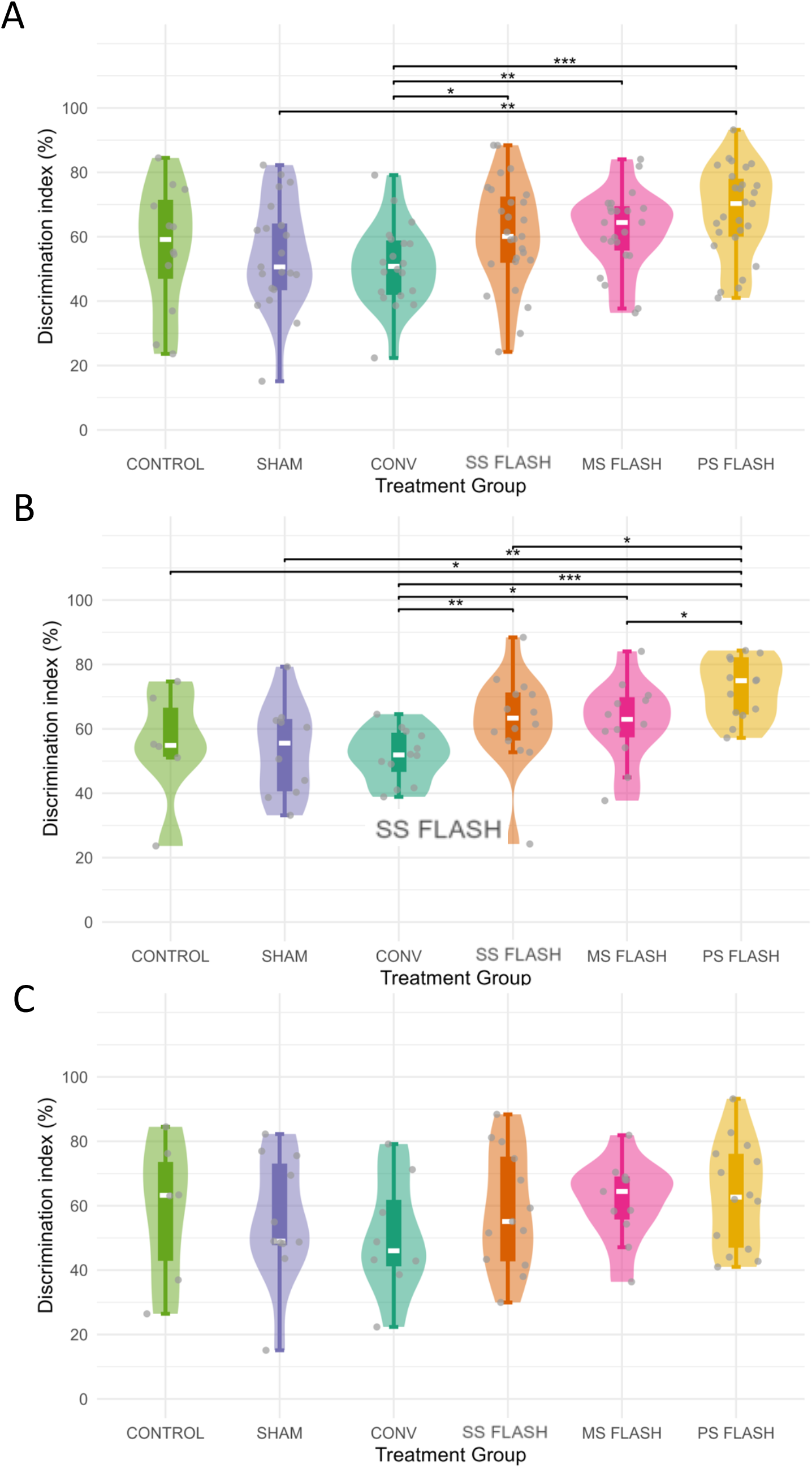
NOR results at 6 weeks post irradiation stratified by treatment group for: (A) all mice (control: n=12, SHAM: n=21, CONV: n=20, SS FLASH: n=27, MS FLASH: n=23, PS FLASH: n=27), (B) male mice (control: n=6, SHAM: n=10, CONV: n=12, SS FLASH: n=14, MS FLASH: n=12, PS FLASH: n=13), and (C) female mice (control: n=6, SHAM: n=11, CONV: n=8, SS FLASH: n=13, MS FLASH: n=11, PS FLASH: n=14). *p<0.05, **p<0.01,***p<0.001. SS: single spill, MS: multi-spill, PS: passive scatter, CONV: conventional

Sex-stratified analyses revealed an important differential effect. In male mice, the pattern mirrored the overall cohort: animals treated with SS, MS, and PS each demonstrated significantly better recognition memory compared to CONV-irradiated males (**Fig. 4B**). Notably, PS-irradiated males demonstrated superior recognition memory relative to all other treatment groups. CONV-irradiated males exhibited the lowest DI (median, [IQR]: 51.90, [10.96]), though this did not differ significantly from SHAM (55.55, [21.24]) or control (54.88, [14.04]; all p>0.05).

In contrast, no significant differences in recognition memory were observed among treatment groups in female mice (**Fig. 4C**). Although CONV-irradiated females demonstrated the numerically lowest median DI (46.00, [19.44]), pairwise comparisons with FLASH-treated, SHAM-irradiated, and unirradiated control females did not reach statistical significance (all p>0.05). The absence of a significant FLASH benefit in females may in part reflect reduced statistical power, as the female CONV subgroup was modestly smaller (n=8) relative to the male subgroup (n=12).

### Visual discrimination and reversal learning touchscreen testing starting at 14 weeks post RT

Visual discrimination and reversal learning were assessed using a touchscreen operant chamber paradigm with training initiated 14 weeks following irradiation. Female mice (SHAM: n=10, CONV: n=11, SS FLASH: n=6) underwent food restriction to 85% of free-feeding body weight prior to and throughout testing to maintain reward salience. No significant difference in body weight was observed across treatment groups at testing initiation. All animals successfully completed pretraining and advanced to visual discrimination.

During visual discrimination, FLASH-irradiated mice completed a significantly greater mean number of trials per session (median, [IQR]: 29.61, [0.36]) compared to both CONV-irradiated (27.85, [2.75]; p=0.024) and SHAM animals (28.44, [4.02]; p=0.022; **Fig.5A**), and demonstrated a significantly shorter mean magazine latency (1.58, [0.44]) relative to CONV-irradiated mice (3.57, [6.52]; p=0.048), suggesting greater reward motivation and task engagement (**Fig. 5C**). No significant differences were observed across groups in mean days required to reach criterion, mean number of correct trials, mean incorrect response latency, or total mean response latency (all p>0.05; **Supplemental Fig. 1**). All animals met visual discrimination criterion and advanced to reversal learning.

**Figure 5.**
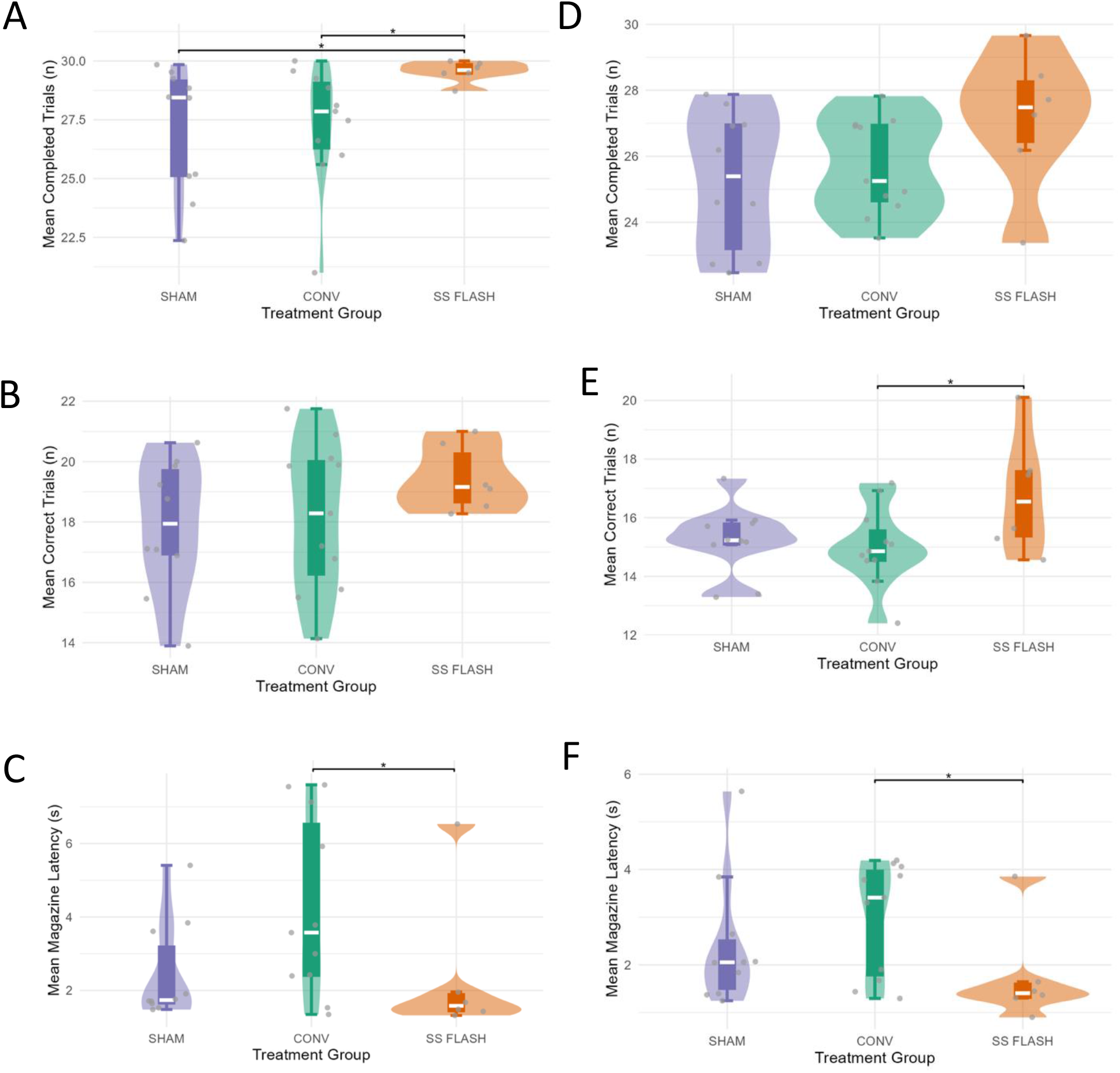
Touchscreen results for visual discrimination across treatment groups (SHAM: n=10, CONV: n=11, SS FLASH: n=6) (A) mean completed trials, (B) mean correct trials, and (C) mean magazine latency. Touchscreen results for reversal learning across treatment groups: (D) mean completed trials, (E) mean correct trials, (F) mean magazine latency. *p<0.05 SS: single spill. CONV: conventional.

During reversal learning, FLASH-irradiated mice demonstrated a significantly greater mean number of correct trials (median, [IQR]: 16.55, [1.01]) compared to CONV-irradiated mice (14.86, [1.01]; p=0.047), with no significant difference observed between FLASH and SHAM animals (25.39, [3.75]; p=0.147), suggesting preservation of cognitive flexibility in FLASH-irradiated mice relative to conventional radiation (**Fig. 5E**). FLASH-irradiated mice again demonstrated a significantly shorter mean magazine latency (1.41, [1.87]) compared to CONV-irradiated animals (3.08, [1.87]; p=0.048; **Fig. 5F**). No significant differences were identified in mean days required to reach criterion, mean completed trials, mean incorrect response latency, or total mean response latency across groups (all p>0.05; **Supplemental Fig. 1**). Reversal learning deficits are considered a preclinical analog of the executive dysfunction and cognitive inflexibility documented in pediatric brain tumor survivors following cranial irradiation, supporting the translational relevance of the observed CONV-specific impairment.

The detection of FLASH-associated cognitive benefits in female mice on touchscreen reversal learning, but not on NOR, may reflect the greater sensitivity of the reversal learning paradigm and the later testing timepoint at which it was administered as the magnitude of RT-associated cognitive deficits increase with time. These findings underscore the importance of multimodal cognitive assessment in preclinical radiation studies and suggest that the neuroprotective profile of FLASH may vary by timepoint.

### MRI at 6 weeks post RT

At 6 weeks post RT, 3 female mice from each treatment group (SS FLASH, CONV and SHAM) underwent MRI. There was no T1 contrast enhancement suggestive of necrosis and no detectable edema on T2 MRI in any mouse. The volume of the lateral ventricles, hippocampus, corpus callosum, external capsule, thalamus, cortex and basal ganglia did not differ among treatment groups (all p>0.05) (**Supplemental Fig. 2B**). Similarly, the mean ADC of the hippocampus, corpus callosum and external capsule also did not differ between treatment groups (**Supplemental Fig. 2C**).

### Histopathology, IF and IHC at 6 weeks post RT

At 6 weeks post-RT, in the hippocampal regions, the percentage of NeuN and GFAP positive cells remained similar across all 3 groups (**Fig. 6A, C**). The percentage of Iba1 positive cells in the CONV group was the highest compared to both FLASH and SHAM groups (9.26 ± 0.64% in CONV vs. 3.91 ± 0.73% in FLASH and 7.00 ± 2.78% in SHAM). A similar trend was observed in the IHC results, where the Iba1 positive area was highest in the CONV group and lowest in the SHAM group (**Fig. 6B, D**). Mean DAB-positive area was 1.65 ± 0.50% tissue in SHAM, 2.66 ± 1.31% in FLASH, and 4.03 ± 0.54% in CONV. Furthermore, the microglia in the CONV group demonstrated prominent soma enlargement with thickened processes and some adopting an amoeboid or rod-shaped morphology. Overall, these features are suggestive of a microglial activation response to CONV, which is largely absent in FLASH.

**Figure 6.**
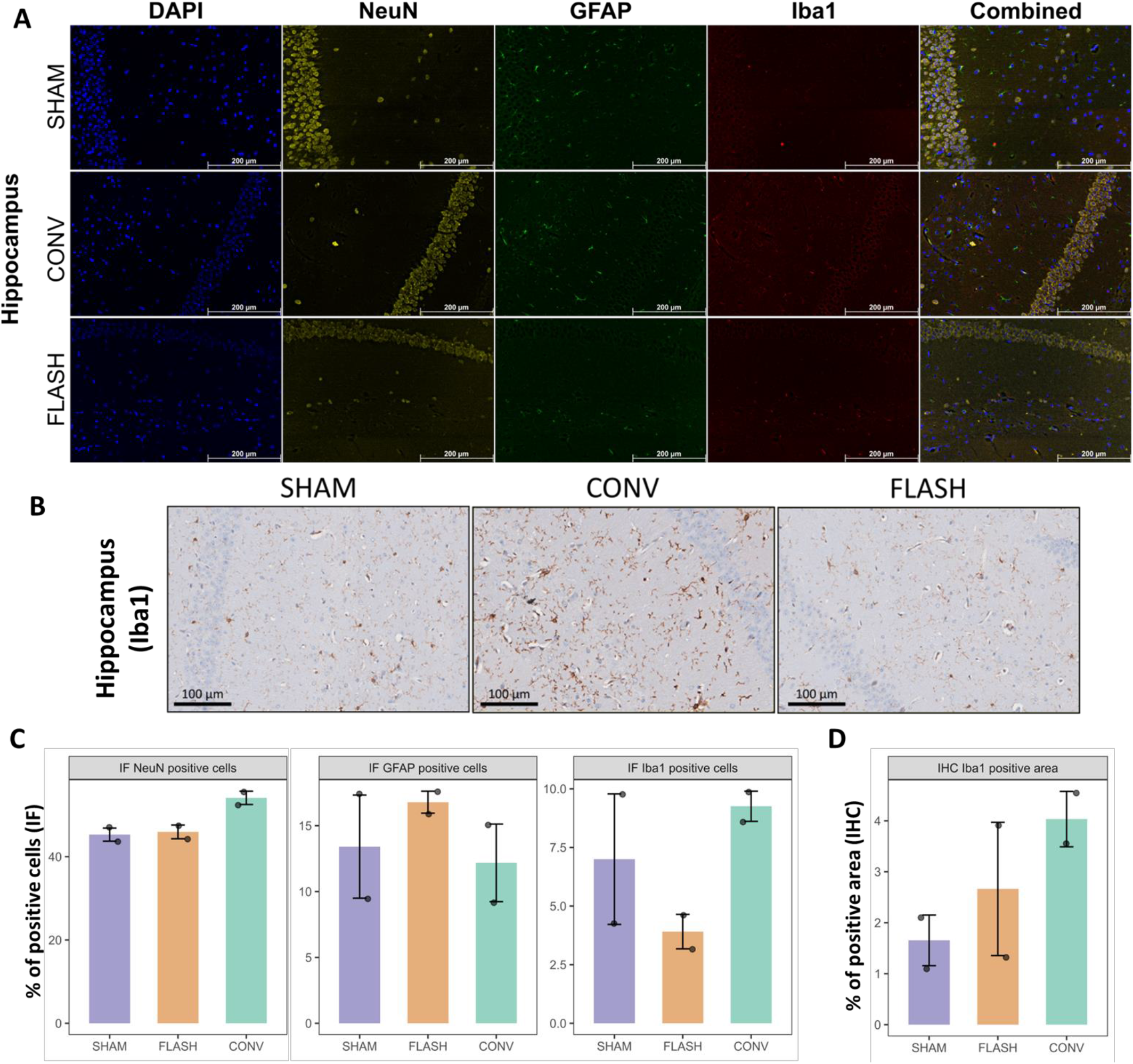
(A) Representative immunofluorescence (IF) imaging of hippocampus on SHAM, CONV and FLASH mice 6 weeks post-RT, with multiplex staining of NeuN (yellow), GFAP (green) and Iba1 (red), alongside DAPI (blue). Scale bar = 200 μm (B) Representative immunohistochemistry (IHC) imaging of hippocampus for Iba1 on SHAM, CONV and FLASH mice. Scale bar = 100 μm (C) Quantification of IF staining. Cells were segmented by DAPI stain and positive cells were identified by positive cytoplasmic staining. (D) Quantification of IHC Iba1 staining through percentage of DAB staining positive area. N=2 for each group (SHAM, FLASH, CONV) across all methods.

## Discussion

Safe clinical translation of proton-FLASH for childhood brain tumors requires defining the delivery conditions under which a neuroprotective effect can be reliably achieved. We demonstrate that FLASH-mediated neuroprotection, as measured by novel object recognition, is preserved across three distinct synchrotron delivery modalities (single-spill, multi-spill, and passive scatter) using a single 10 Gy whole-brain dose. This is particularly relevant for clinical translation, as the charge deliverable in a single synchrotron spill is inherently limited, and multi-spill delivery strategies will likely be necessary to achieve target coverage in human-scale volumes. Critically, FLASH conditions (≥40 Gy/s) were met across all three modalities only when dose rate was quantified using the dose-averaged dose rate (DADR) framework; conventional metrics such as PBS dose rate and field dose rate failed to classify these deliveries as FLASH-qualifying, despite the observed neuroprotective effect in all three groups. Unlike field dose rate, which averages over the entire delivery window, or instantaneous metrics that reflect only individual spot delivery, DADR weights each contributing dose element by its fractional contribution to the total dose, more accurately capturing the effective dose rate experienced by a given tissue volume across the full temporal structure of beam delivery. This dissociation supports DADR as the most biologically relevant dose rate metric for synchrotron-based PBS delivery defined to-date, and suggests that multi-spill proton-FLASH may be a viable clinical strategy for large target volumes provided that per-spill dose rates exceed 40 Gy/s.

The lack of standardization in dose rate reporting for PBS delivery has been a recognized challenge in the field. Folkerts et al. introduced the PBS dose rate metric, defining effective irradiation time using a dose threshold to account for the spatiotemporal structure of spot delivery,^12^ while van de Water et al. proposed DADR as a dose-weighted instantaneous metric that instead disregards inter-spot intervals.^13^ Subsequent comparative analyses have demonstrated that these definitions yield substantially different values for the same delivery: comparison of five metrics applied to a single-energy head and neck plan demonstrated that DADR values were approximately twice those of PBS-based metrics, because DADR excludes periods when a voxel is not actively receiving dose.^26^ There is also a large variation in OAR dose rate classification depending on how dose rate was calculated for PBS transmission beam plans, concluding that the choice of metric poses direct challenges to FLASH clinical implementation and trial design.^27^ However, none of these prior studies could resolve which metric best predicts biological outcome, as they were dosimetric analyses without paired in vivo endpoints. Our data provide the first biological evidence to inform this question: by demonstrating neuroprotection across single-spill, multi-spill, and passive scatter deliveries that met UHDR thresholds only by DADR, we show that DADR is not merely a more permissive calculation but may more accurately reflect the dose rate parameter driving the FLASH effect in brain tissue — consistent with the interpretation that inter-spot temporal gaps are biologically less relevant than the instantaneous rate experienced during active dose delivery.

The published data on proton FLASH neuroprotection are mixed and must be interpreted in the context of dose, species, beam modality, and the cognitive domains assessed. In neonatal rats, FLASH-mediated sparing was observed at 5 Gy but not 8 Gy whole brain irradiation,^28^ and in adult rats, an 18 Gy single fraction failed to produce neuroprotection across a broad behavioral battery.^11^ Since electron FLASH has been shown to confer neuroprotection in juvenile mice at 10 Gy,^29^ developmental stage alone does not account for the absence of proton FLASH sparing at 8 Gy in the neonatal rat — pointing instead to species-specific differences in radiosensitivity, potential differences in the biological effectiveness of protons relative to electrons at equivalent doses, or the possibility that the specific cognitive domains and behavioral paradigms tested in rat models are less sensitive to detecting partial neuroprotection than the NOR task used in murine studies. Our results add to this literature by demonstrating a neuroprotective proton FLASH effect at 10 Gy whole brain irradiation in adult mice, consistent with the dose at which electron FLASH was originally shown to spare memory. We included an equal distribution of male and female mice, allowing us to detect sex-dependent patterns that have been largely unexamined in the proton FLASH literature. While males demonstrated a greater memory benefit on NOR at 6 weeks, females showed superior cognitive flexibility and processing speed on touchscreen training and reversal learning, which was initiated at 14 weeks post-irradiation. The detection of a FLASH-associated benefit in females on reversal learning, but not on NOR, most likely reflects both the greater sensitivity of the reversal learning paradigm and the later timepoint at which it was administered, as the magnitude of radiation-associated cognitive deficits is known to increase progressively with time after cranial irradiation. Notably, reversal learning deficits are widely considered a preclinical analog of the executive dysfunction and cognitive inflexibility documented in pediatric brain tumor survivors following cranial irradiation, lending direct translational relevance to the observed conventional dose rate-specific impairment in this domain. Taken together, these findings underscore the importance of multimodal cognitive assessment across multiple timepoints and both sexes in preclinical radiation studies, as reliance on a single endpoint or a single sex risks underestimating both the burden of radiation-induced cognitive injury and the potential benefits of FLASH delivery.

Cranial irradiation-induced neurocognitive impairment in children is associated with progressive structural brain changes, particularly reductions in hippocampal and corpus callosum volume over time, with the most vulnerable domains being memory and processing speed.^22^ Complementing these volumetric findings, diffusion-weighted imaging studies have demonstrated increasing hippocampal mean diffusivity and decreasing corpus callosum, trajectories that are opposite to what is observed in healthy children and irradiated survivors with normal neurocognition.^23^ Although we obtained MRI at 6 weeks post RT, we did not observe changes in volume or ADC between FLASH and CONV treatment groups. This null finding is likely attributable to several factors: the early imaging timepoint may precede the emergence of detectable structural divergence between dose rate groups, the in vivo acquisition parameters — including resolution and slice thickness — may have been insufficient to capture subtle intergroup differences in a small animal model, and the sensitivity of ADC as a single diffusion metric is inherently limited compared to more comprehensive diffusion tensor imaging approaches. Ex vivo imaging offers a promising path forward, as it enables substantially higher spatial resolution, acquisition of a greater number of diffusion directions, and full diffusion tensor modeling — capabilities that may reveal microstructural differences between FLASH and CONV that were below the detection threshold of our in vivo protocol and that may ultimately link the neuroprotective behavioral phenotype we observed to an underlying structural substrate.

Radiation-induced cognitive dysfunction is linked to neuroinflammation, oxidative stress and a decrease in neurogenesis, as well as disruptions to neuronal synapses.^30^ The interplay between neurons, astrocytes, and microglia is crucial in understanding radiation-induced neuroinflammatory responses. The release of various cytokines and chemokines after radiation causes neurons to secrete high mobility group box-1 (HMGB1) that bind to Toll-like receptor 4 (TLR4) and leads to a downstream cascade that activates microglia.^31^ Astrocytes, in response to radiation, can also be induced to a neurotoxic state that further secretes pro-inflammatory cytokines.^32^ Consequently, this further induces the microglia to an active state. While there are many proposed pathways by which radiation induces microglial activation and subsequent neuroinflammation, one proposed mechanism of action involves the activation of the cyclic GMP-AMP synthase (cGAS)-stimulator of interferon genes (STING) signaling pathway after radiation that mediates persistent microglia activation and neuroinflammation, leading to neural degeneration.^33^ In a previous study, radiation was shown to trigger this activation via the accumulation of reactive oxygen species (ROS) that caused leakage of mitochondrial DNA into the cytoplasm, activating cGAS-STING mediated microglial activation. In the context of FLASH irradiation, a study following mice 6 months post electron RT found that microglia activation (Iba1/CD68) was significantly higher in the CONV group as compared to FLASH, in line with the decreased neuroinflammation and better neurocognitive outcomes observed in the FLASH group.^6^

Our results further validate existing findings in a proton-specific context, where we see similar trends in Iba1 presence that could be linked to the neurocognitive outcomes we observed. While our IF and IHC results were limited to 2 mice per group, we observed the expected trend of a lower number of Iba1 positivity through both IF and IHC in the FLASH compared to CONV group. Although we are unable to distinguish the exact homeostatic or activated state of the microglia through Iba1 staining alone, the reduced presence of these cells and lack of activated-state morphology on IHC could indicate a decrease in neuroinflammation in the FLASH mice. This possibly contributes to the improved neurocognitive outcomes we observed.

Several limitations of this study warrant consideration. First, neurocognitive outcomes were assessed following a single 10 Gy whole-brain dose; the dose-rate dependence of the FLASH effect across fractionated regimens and craniospinal delivery — the conditions under which this technology will most plausibly enter pediatric clinical use — remains to be defined and is the subject of ongoing work. Second, our microglial analysis was limited to two mice per group and used Iba1 as the sole marker; definitive characterization of microglial state requires combined assessment of homeostatic and activation markers and formal morphometric quantification, which is the subject of follow-up single-cell transcriptomic work. Third, MRI was acquired at a single timepoint 6 weeks post-RT, which may precede the structural changes that emerge progressively in pediatric brain tumor survivors, and ex vivo high-resolution diffusion imaging may be required to detect microstructural differences below the detection threshold of our in vivo protocol. Finally, touchscreen-based assessment was performed only in female mice due to logistical constraints, limiting our ability to evaluate sex-by-modality interactions on this paradigm. Despite these limitations, the convergent evidence from recognition memory, cognitive flexibility, and microglial endpoints supports the central finding that synchrotron-based proton FLASH preserves neurocognitive function across delivery modalities relevant to clinical translation.

## Conclusion

In this study, we demonstrate that the neuroprotective FLASH effect is preserved across three synchrotron-based proton delivery modalities — single-spill PBS, multi-spill PBS, and passive scatter — using a single 10 Gy whole-brain dose. Critically, the FLASH effect was maintained under multi-spill delivery, where conventional dose-rate metrics (FDR, PBSDR) fall below the 40 Gy/s threshold and only the dose-averaged dose rate (DADR) qualifies the delivery as FLASH. This dissociation between dosimetric classification and biological outcome supports DADR as the dose-rate metric most relevant to the FLASH effect in brain tissue and provides direct biological evidence informing a long-standing question in the field. The preservation of neuroprotection under multi-spill delivery is essential for clinical translation, as larger human target volumes — particularly in pediatric brain tumors requiring whole-brain or craniospinal coverage — cannot be encompassed within a single proton spill on synchrotron-based systems. Multimodal cognitive assessment revealed sex- and timepoint-dependent patterns, with males showing FLASH-mediated preservation of recognition memory at 6 weeks and females showing preservation of cognitive flexibility on reversal learning at later timepoints, underscoring that single-domain or single-timepoint assessment risks underestimating the FLASH effect. Reduced Iba1^+^ presence in FLASH-treated mice supports a microglial contribution to the neuroprotective mechanism. Together, these findings establish the feasibility and biological rationale for synchrotron-based proton FLASH as a strategy to reduce treatment-related neurocognitive impairment in children with brain tumors, and provide a framework for the dose-rate reporting standards needed to support clinical trial design.

## Acknowledgements

The authors thank Shibuya Shinji and Paul Boisseau for their invaluable support during the proton UHDR experiments. We thank Mohammed Rezaee for his assistance with initial concept development and feedback throughout the project.

