## Supplementary material for "Proton FLASH preserves neurocognition across delivery techniques: implications for clinical translation in pediatric brain tumors": Supplmental Figures

Supplemental Figure 1

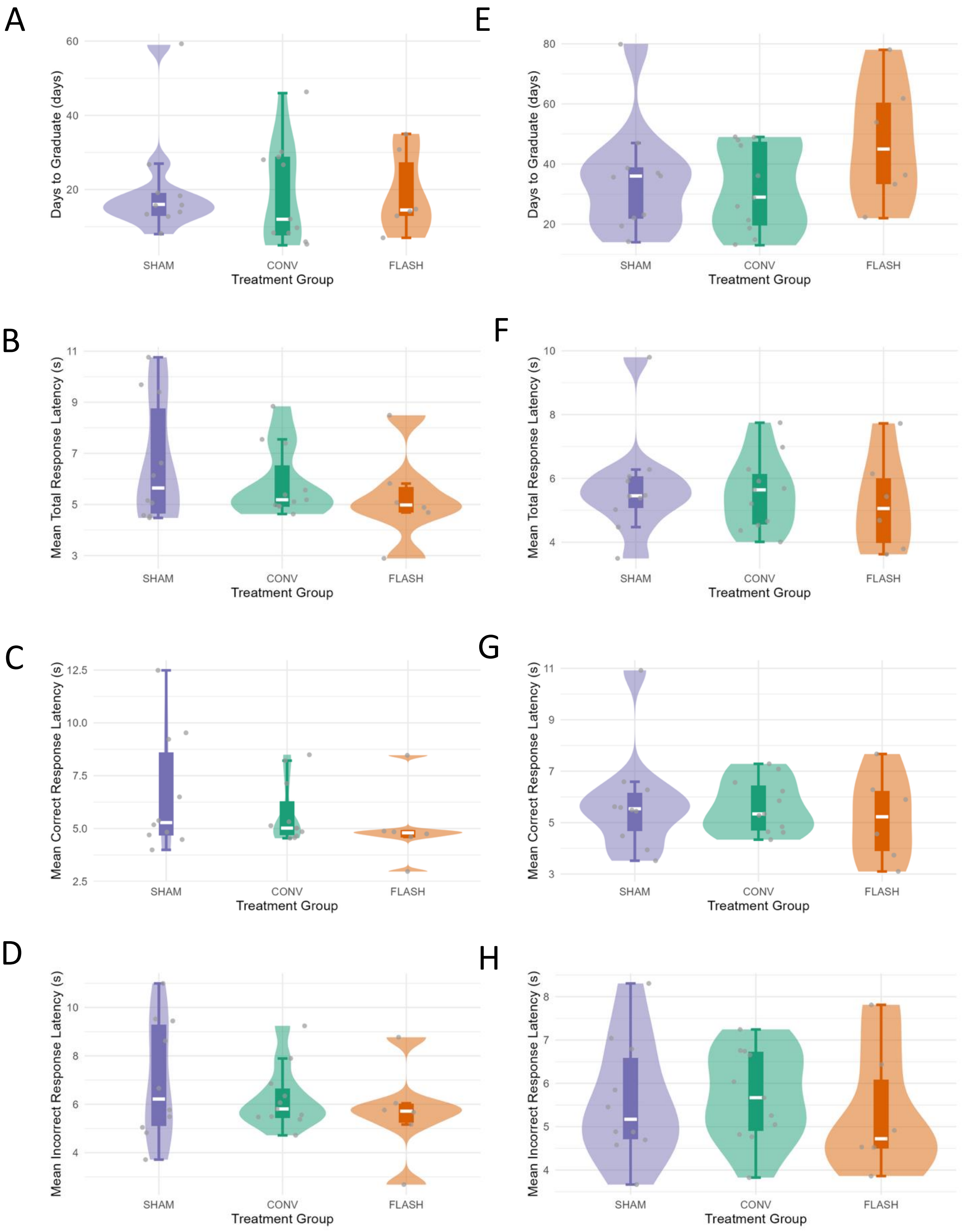


**Supplemental Figure 1**. Touchscreen results for visual discrimination across treatment groups (SHAM: n=10, CONV: n=11, SS FLASH: n=6): (A) days to graduate, (B) mean total response latency, and (C) mean correct response latency, (D) mean incorrect response latency. Touchscreen results for reversal learning across treatment groups: (A) days to graduate, (B) mean total response latency, and (C) mean correct response latency, (D) mean incorrect response latency. *$p<0.05$ SS: single spill. CONV: conventional.

Supplemental Figure 2

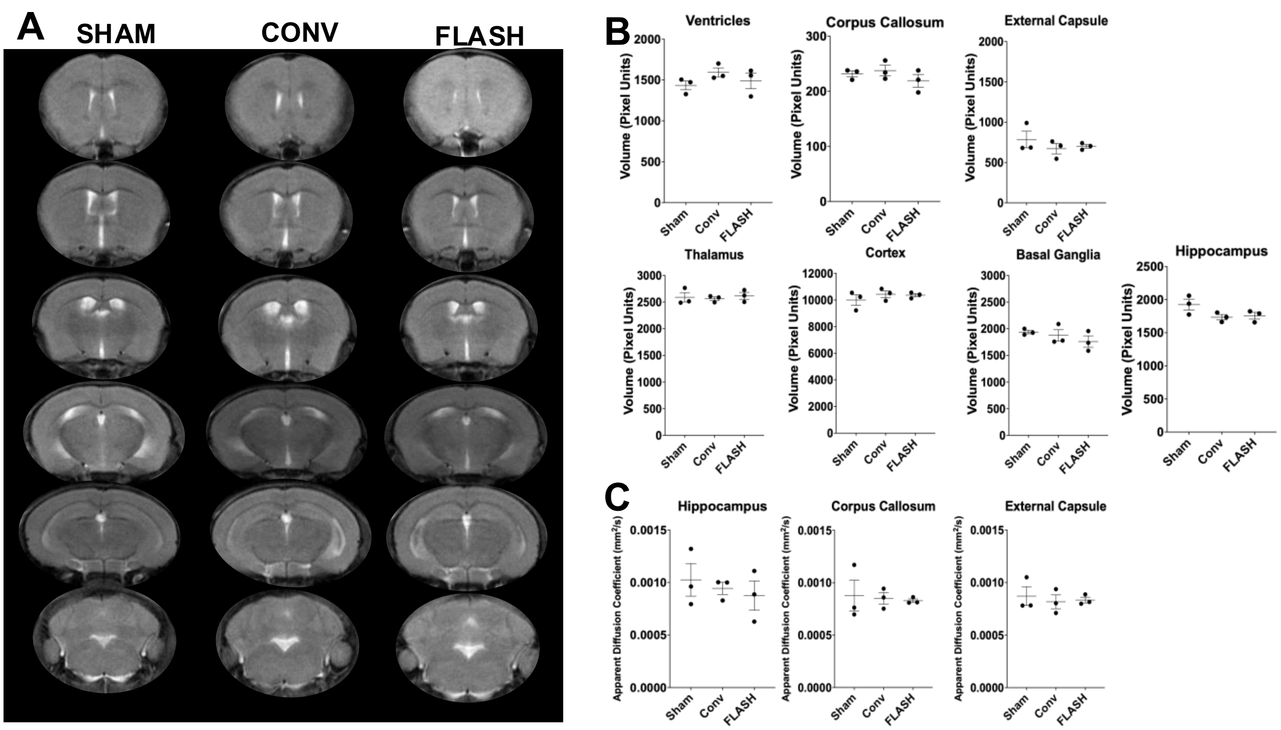


**Supplemental Figure 2**. MRI was acquired 6 weeks post irradiation in 3 mice per treatment group (SHAM, CONV, SS FLASH). (A) Representative coronal images on T2 MRI stratified by treatment group. (B) Volume of brain substructures by treatment group. (C) ADC by treatment group. Median and interquartile range shown for each scatter plot. ADC: apparent diffusion coefficient.
